# Mathematical model for visual cortical development based on molecular mechanisms underlying the generation and elimination of dendritic spines

**DOI:** 10.64898/2026.10.24.761552

**Authors:** Shigeru Tanaka, Masanobu Miyashita

## Abstract

In the developing brain, neural circuits are elaborated by the activity-dependent rewiring of synaptic connections. The current view of synaptic rewiring in early development is that filopodia spontaneously generated along dendrites transform into spines by establishing contacts with neighboring axonal boutons to form synapses whereas spines of inappropriate synapses are eliminated in an activity-dependent manner. In this theoretical study, we mainly focus on the processes of spine generation, maturation, and elimination to build a mathematical model of synaptic rewiring, incorporating experimental findings on actin polymerization and depolymerization under the activation of Rho-GTPases mediated by the binding of brain-derived neurotrophic factor (BDNF) to neurotrophin receptors such as tropomyosin kinase receptor B (TrkB) and the low-affinity-nerve growth factor receptor (p75^NTR^). TrkB receptors are dephosphorylated by chondroitin sulphate proteoglycans (CSPGs) mediated by protein-tyrosine phosphatase σ (PTPσ) in an age-dependent manner. The model describes the generation or elimination of dendritic spines of visual cortical neurons under actin polymerization and depolymerization to establish appropriate synaptic contacts on afferent fibers of cells in the lateral geniculate nucleus. The simulations based on the model reproduced not only simple-cell-like receptive fields and orientation-direction maps but also a characteristic profile of the total number of synapses, which shows an early increase and a delayed decrease during the critical period. It is also demonstrated that the formation of direction selectivity is delayed behind the formation of orientation selectivity, which is likely promoted by the pruning of inappropriate synapses.

## 1. Introduction

In early development, abundant membranous protrusions, filopodia, emerge spontaneously on dendrites, and then transform into spines in an activity-dependent manner (**Jontes and Smith, 2000**). Such structural plasticity of spines is attributed to the dynamic changes in cytoskeletal actin filaments. Here we pay special attention to the brain-derived neurotrophic factor (BDNF) (**Binder and Scharfman, 2004; Cunha et al., 2010**) and some related molecules to describe the activity-dependent transformation of spines from filopodia and the elimination of spines. Among Rho-GTPases, RhoA, Cdc42, and Rac1 control the dynamics of actin cytoskeletons of dendritic spines (**Ridley, 2006; Murakoshi et al., 2011; Sala and Segal, 2014; Hedrick et al., 2016**). The activations of Cdc42 and Rac1 (**Murakoshi et al., 2011; Saneyoshi and Hayashi, 2012; Hedrick et al., 2016**) through the activation of tropomyosin kinase receptor B (TrkB) (**Klein et al., 1989**) facilitate actin polymerization, resulting in spine growth, whereas RhoA contributes to the shrinkage or collapse of spines through the activation of the low-affinity nerve growth factor receptor (p75^NTR^) (**Singh et al., 2008; Je et al., 2012; Sun et al., 2012**). proBDNF and mature BDNF are stored inside dense core vesicles in cortical neurons and released via two pathways: the regulatory pathway and the constitutive pathway through exocytosis (**Thomas and Davies, 2005**). The secretion of BDNF via the regulatory pathway takes place depending on the intracellular Ca^2+^ concentration (**Kolarow et al., 2007**). Mature BDNF and proBDNF released from a dendritic spine via either pathway bind to TrkB and p75^NTR^ expressed on the spine surface, respectively. Furthermore, tissue-plasminogen activator (tPA) (**Tsai, 2017**) is released in an activity-dependent manner, and the resultant plasmin cleaves proBDNF into mature BDNF and BDNF propeptide in the extracellular space (**Greenberg et al., 2009; Nagappan et al., 2009**). On the other hand, the dephosphorylation of TrkB by protein-tyrosine phosphatase σ (PTPσ) (**Kurihara and Yamashita, 2012**), which is a receptor of chondroitin sulfate proteoglycans (CSPGs) (**de Vivo et al., 2013**), inhibits the function of TrkB. In parallel, CSPGs, which are major components of the extracellular matrix in large amounts, act as physical obstacles (**Levy et al., 2014**) that reduce spine motility, regulating the probability that a filopodia or spine encounters a presynaptic axonal bouton.

Considering the above-mentioned molecular mechanisms, we derive the probability that one spine is added to or eliminated from a dendrite. In computer simulations, synapses are rewired through the generation and elimination of spines to change the connectivity of afferent inputs to cortical neurons. Through these processes, afferent inputs compete with each other to form synapses at cortical neurons and, finally, simple-cell-like spatiotemporal receptive fields and orientation-direction maps are formed, as observed in cat area 17 (**DeAngelis et al., 1995; Kim et al., 1999; Swindale et al., 1987**). We assume that the amount of p75^NTR^ in the visual cortex decreases depending on light exposure around eye opening (**Bracken and Turrigiano, 2009**), and the amount of TrkB is decreased by the dephosphorylation of TrkB mediated by CSPGs via PTPσ in the late stage of the critical period (**Kurihara and Yamashita, 2012**), as observed in experimental studies (**Grigonis et al., 1988; Bourgeois and Rakic, 1993; Elston et al., 2009; Faust et al., 2021**). We find that the number of synapses increases initially, then decreases asymptotically and becomes a constant number towards the end of the critical period. In addition, orientation selectivity almost monotonically increases towards maximum selectivity until the end of the critical period. In contrast, direction selectivity increases later than orientation selectivity, which suggests that direction selectivity formation is associated with the pruning of inappropriate synapses. We consider that the closure of the critical period is caused by the prolongation of time for spines to find candidate axonal boutons to form synaptic contacts, which may be accompanied by reduced spine motility.

## 2. Materials and Methods

We built a mathematical model of the developmental rewiring of afferent inputs from lateral geniculate cells to neurons in layer 4 of the visual cortex on the basis of the following molecular mechanisms.

1. The intracellular Ca^2+^ concentration in the visual cortical neurons is increased by Ca^2+^ influx through the N-methyl-D-aspartate receptor (NMDAR) activated by visual stimulation and is enhanced by the occurrence of backpropagating action potentials (**Koester and Sakmann, 1998; 2000**).
2. The exocytosis of mature BDNF and proBDNF from dense core vesicles occurs in a Ca^2+^ -concentration-dependent manner in the regulatory pathway and in an activity-independent manner in the constitutive pathway.
3. Tissue plasminogen activator (tPA) is also released in an activity-dependent manner, which converts plasminogen into plasmin (**Pang et al., 2004; Lochner et al., 2005**). Plasmin cleaves proBDNF into mature BDNF and BDNF pro-peptide in an extracellular space.
4. Mature BDNF binds TrkB expressed on the spine surface, which activates Cdc42 and Rac1.
5. The activated Cdc42 and Rac1 promote actin polymerization, resulting in the elongation of spine necks and the enlargement of spine heads (**Govek et al., 2005**).
6. BDNF pro-peptide and proBDNF bind to the pan-neurotrophin receptor (p75^NTR^), which activates RhoA (**Sun et al., 2012**).
7. The activated RhoA diffusing in the dendrite depolymerizes actin filaments, which leads to the shrinkage of adjacent dendritic spines (**Koo et al., 2015**).
8. Upon eye opening, the expression level of p75^NTR^ starts to decrease, which is likely to contribute to the increase in the number of synapses.
9. As TrkB is dephosphorylated by PTP*σ* activated by CSPGs, the amount of TrkB available for mature BDNF gradually decreases and the number of synapses also decreases (**Kurihara and Yamashita, 2012**).
10. The expression of CSPGs itself reduces the motility of dendritic spines through the formation of a linkage between spines and axonal boutons via cell adhesion molecules such as integrins and the cadherin/catenin complex (**Nelson, 2008**), and, finally, the critical period closes.

A mathematical model is built on the basis of the above-mentioned molecular mechanisms, which control actin polymerization and depolymerization that change the dendritic spine morphology and finally synapse formation and elimination in an activity-dependent manner. The activity-dependent alteration in dendritic spines through the competitive interactions among nearby spines is described by the Lotka-Volterra equation (**Hofbauer and Sigmund, 1988; Fukai and Tanaka, 1997**). This equation may describe rapid volume changes in dendritic spines in living slice preparations under local stimulation by glutamate (**Callaway and Katz, 1993**) or BDNF (**Kossel et al., 2001**). However, in this paper, we focus on the developmental changes in the synaptic number and response selectivity in visual cortical neurons.

### 2.1 Neuronal activities

The membrane potential of the *i* th visual cortical neuron 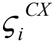 is given as the sum of the excitatory postsynaptic potential evoked by the firing of LGN cells, the inhibitory potential induced by the indirect geniculate inputs via weakly oriented interneurons in layer 4 (**Miller, 2003**), and the membrane potential evoked by other cortical neurons through excitatory and inhibitory lateral connections is expressed as

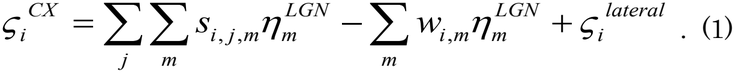

Here, 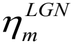 represents the firing rate of the *m* th LGN cell, and *j* is the index of a filopodium or spine along the dendrite. *s*_*i, j*,*m*_ represents the synaptic strength between the *j* th dendritic spine site of the *i* th cortical neuron and the *m* th LGN cell. The receptive field of an LGN cell is indicated by (*k, μ*_1_, *μ*_2_), where *k*, *μ*_1_, and *μ*_2_ represent the retinotopic center position, ON- or OFF-center type, and nonlagged or lagged type of LGN cell, respectively. Here, if we assume that the excitation and inhibition induced by feedforward inputs from LGN cells are balanced on average, 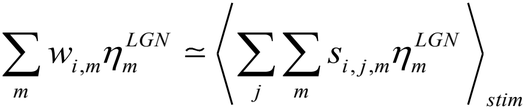. 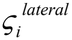 is written as

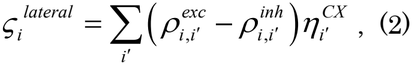

where 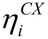 represents the *i* th cortical neuron’s firing rate. 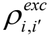 and 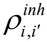 represent excitatory and inhibitory lateral connections, respectively. In this study, we do not consider the changes in the intracortical connectivity, assuming that the maturation of lateral connections occurs earlier than that of afferent inputs. Lateral inhibition signals via inhibitory interneurons polysynaptically, whereas lateral excitation signals are conveyed by monosynaptic connections from neighboring excitatory neurons. This suggests that the range of lateral inhibition is likely wider than that of lateral excitation. Thus, 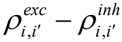 is written as the Mexican-hat-type cortical interaction function 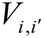 of the cortical distance 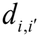 between neurons *i* and *i*′ :

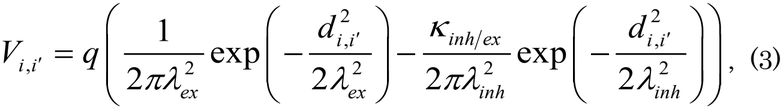

where the parameters *q, K*_*inh ex*_, *λex*, and *λ*_*inh*_ represent the interaction strength, the ratio in strength of inhibition to excitation, the spatial extent of excitatory connections, and the extent of inhibitory connections, respectively.

The membrane depolarization is assumed to be given by the half-rectification of the total membrane potential 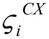 (**Cash and Yuste, 1998**) with the threshold 0:

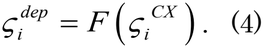

The firing rate 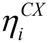 is also assumed to be given by the rectification of the membrane depolarization with the firing threshold *ϑ* > 0 . That is,

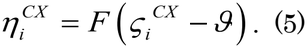

Using the Heaviside function Θ (*x*) : Θ(*x*) = 1 for *x* > 0, 0 otherwise, we obtain the rectification function *F* (*x*) as

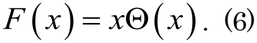

Here, it is assumed to be set as 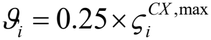, where 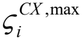 represents the maximum value of 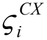.

### 2.2 Activity-dependent calcium influx

Calcium ions flow into dendritic spines from the extracellular space through NMDA channels on the spine surface when the NMDAR is activated while the postsynaptic potential is depolarized. Then, the intracellular calcium concentration at the *j* th spine is further increased by the occurrence of the backpropagating action potential (bAP) of a cortical neuron (**Markram et al., 1997; Koester and Sakmann, 1998; Koester and Sakmann, 2000**). The intracellular calcium concentration at the *j* th spine is modeled as the sum of these effects (**Nevian and Sakmann, 2004; Jia et al., 2014**):

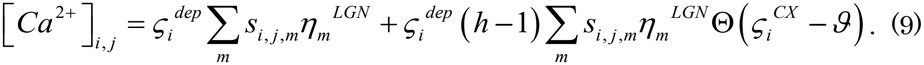

The first term on the right-hand side represents calcium concentration taken up by the NMDA channel activation associated with the LGN cell’s firing, and the second term indicates calcium concentration elevated by the bAP, where the Heaviside function indicates the firing event occurring at the *i* th neuron. *h* (> 1) is the calcium concentration enhancement factor.

Using the spine volume *ρi, j* at the *j* th spine of the *i* th neuron, and the afferent inputs from the *m* th LGN cell to the *j* th spine*σ* _*i, j*,*m*_, we obtain the synaptic strength *s*_*i, j*,*m*_ as

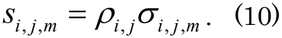

Therefore,

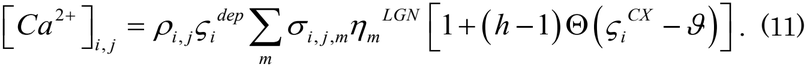

It is widely accepted that a single spine forms synaptic contact with a single axonal bouton. Thus, if the *j* th spine on the dendrite of the *i* th cortical neuron makes contact with a bouton of the axon from the *m* th cell of the LGN, *σ* _*i, j*,*m*_ = 1, and if there is no contact between the *j* th spine and the *m* th cell *σ*_*i, j*,*m*_ = 0.

### 2.3 BDNF release and neurotrophin receptor activation

BDNF is stored within dense core vesicles in cortical neurons as a mixture of mature BDNF (mBDNF) and proBDNF, where proBDNF is observed to be more abundant than mBDNF (**Nagappan et al., 2009**). The ratios of mBDNF to proBDNF within vesicles in the regulatory and constitutive pathways are set to be *ω* to 1 − *ω* and *ω*′ to 1 − *ω*′, respectively. The secreted mBDNF and proBDNF are given by

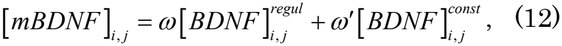

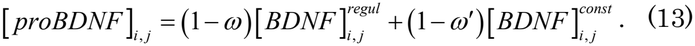

The total amount of BDNF secreted from the regulatory pathway 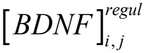 is proportional to the intracellular calcium concentration [*Ca*^2+^]_*i, j*_ and given by [*Ca*^2+^]_*i, j*_ using the proportional constant *K* . In contrast, the amount of BDNF released from the constitutive pathway, 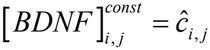, may be probabilistically different at different dendritic positions, where the secreted amount of BDNF 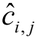 is a stochastic variable. Thus, we obtain

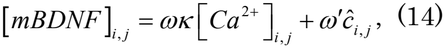

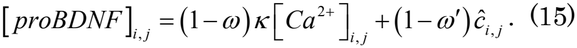

It has been reported that proBDNF released from the regulatory pathway is cleaved into mBDNF and BDNF propeptide by the plasmin generated from plasminogen converted by tPA released from cortical neurons in an activity-dependent manner. Considering the complete conversion of proBDNF into mBDNF and BDNF propeptide by the extracellular plasmin, the extracellular concentration of the three variants of BDNF is given as

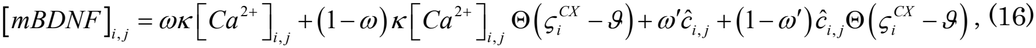

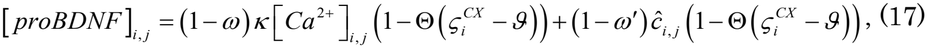

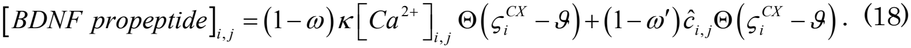

Since the BDNF propeptide exhibits high affinity to p75^NTR^ as proBDNF does, we can sum up [*proBDNF*]_*i*, *j*_ and [*BDNF propeptide*]_*i*, *j*_ to get

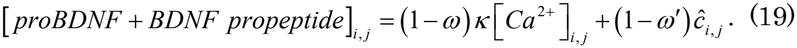

When we assume that the amount of activated TrkB, 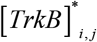, at the *j* the dendritic position of the *i* th neuron is proportional to the product of [*mBDNF*] _*i, j*_ and [*TrkB*], where for simplicity we assume that TrkB is uniformly expressed on the dendrites of visual cortical neurons,

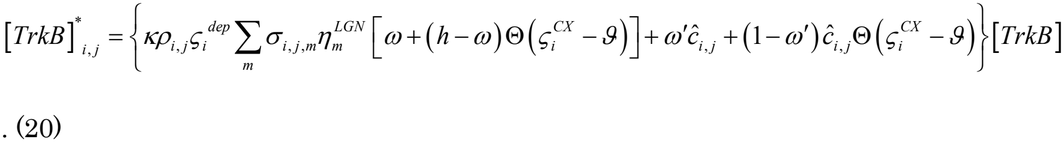

Here, we use the property of the Heaviside function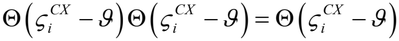 . Likewise, the amount of activated p75^NTR^,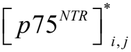, at the *j* th dendritic position of the *i* th neuron is proportional to the product of [*proBDNF* ^+^ *BDNFpropeptide*]_*i, j*_ and [*p*75^*NTR*^] .

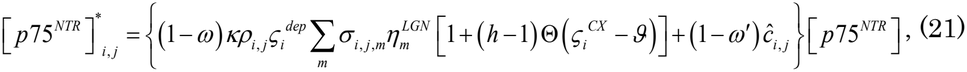

where p75^NTR^ is assumed to be uniformly expressed on the dendrites in the visual cortical neurons

### 2.4 Spine dynamics assisted by Rho-GTPases

Actin polymerization and depolymerization are regulated by Rho-GTPases such as Cdc42, RhoA, and Rac1. Cdc42 and RhoA are related to the elongation and shrinkage of actin filaments, respectively. On the other hand, Rac1 contributes to the branching and networking of actin filaments inside spines, which result in the expansion of the areas of spine heads. Rac1 may be more involved in the induction of LTP than in the morphological changes of dendritic spines related to synaptic rewiring. In this study, since we are more interested in the modeling of spine formation and elimination, we derive the dynamic equation for dendritic spine elongation and shrinkage. Here, we identify a locally accumulated amount of actin filaments as a spine volume. For simplicity, we disregard the effect of the diffusion of Rac1 in the derivation of the dynamic equation of a spine volume:

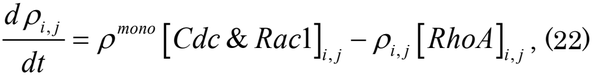

where *ρ* ^*mono*^ represents the amount of supplied actin monomers (G-actin) inside individual spines. The first term on the right-hand side of Eq. (22) indicates the rate of polymerization of actin monomers assisted by Cdc42 and Rac1, and the second term indicates the rate of collapse of actin filaments under the effect of RhoA.

On the other hand, RhoA spreads diffusely from spines to dendritic segments with the tendency to shrink adjacent spines. Rac1 is also diffused to adjacent spines, branching actin filaments and forming mushroom spines, whereas Cdc42 is localized within individual spines and elongates spine necks. For simplicity, we neglect the diffusivity of Rac1 here: the diffusion coefficients of Cdc42 and Rac1 are set to 0, and their lifetimes are commonly set to be *τ* _*GTP*_ . The product [*RhoA*]_*i, j*_ *ρ*_*i*, *j*_ represents the rate of depolymerization of actin filaments, leading to the pruning of the *j* th spine from the ith neuron.

Let us assume that *D*_*rhoA*_ and *τ* _*GTP*_ are the diffusion coefficient and lifetime of activated RhoA, respectively. Equations of the concentrations of activated Cdc42 & Rac1 and RhoA are respectively given by

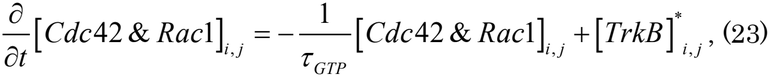

and

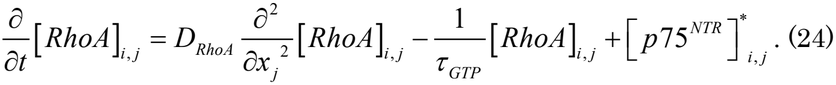

Because the dynamics of Rho-GTPases is a process sufficiently faster than the generation and elimination of single spines, it is considered that [*Cdc*42 & *Rac*1]_*i, j*_ and [*RhoA*]_*i, j*_ reach rapidly the steady-state concentrations given as solutions to Eqs. (23) and (24):

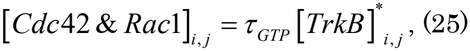

and

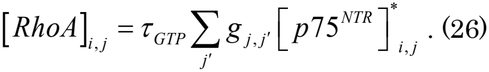

The diffusion kernel 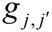 in Eq. (26) is given by

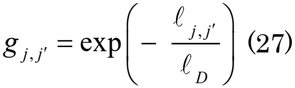

with the distance 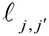 between a pair of spines at *j* and *j*′ on a single dendrite and the diffusion length of RhoA,

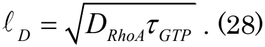

Now, we can rewrite Eq. (22) as

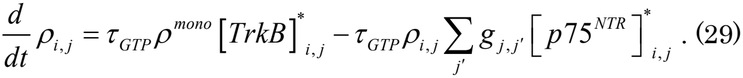

If the BDNF in the constitutive pathway is also released at spine sites, 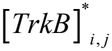 and 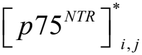 can be rewritten respectively as

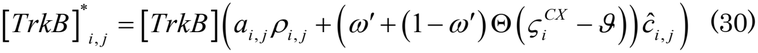

and

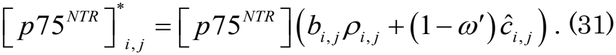

Now, we introduce 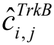 and 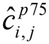 as

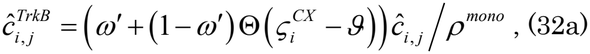

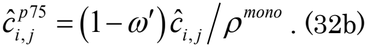

When we discuss synaptic rewiring during development, the calcium concentration should be time-averaged within the time required for the generation/elimination of synapses. Thus, we introduce *a*_*i, j*_ and *b*_*i, j*_, which are given by

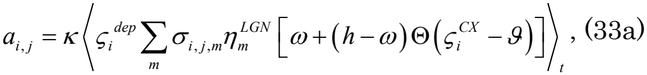

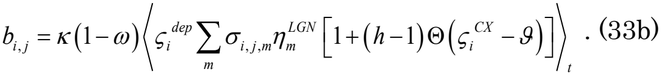

Equation (29) is given by

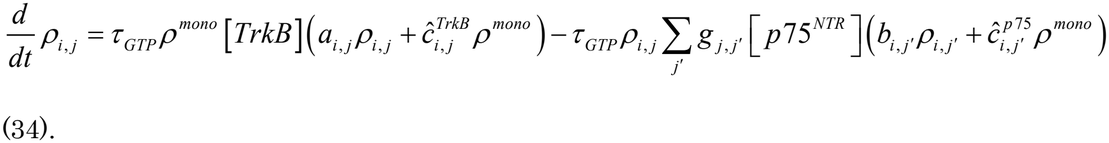

This leads to the dynamic equation of the normalized spine volume 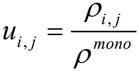 with the rescaled time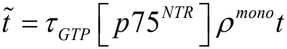 as a multivariable competitive Lotka-Volterra-type equation (**Hofbauer and Sigmund, 1988; Fukai and Tanaka, 1997**):

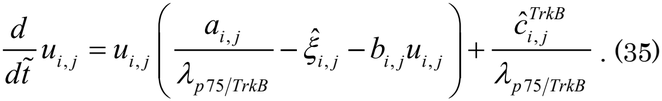

Note that *λ*_*p* 75/*TrkB*_ and 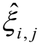are respectively defined by

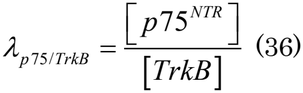

and

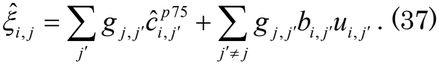

The amount of BDNF secreted from the constitutive pathway is expected to be sufficiently smaller than that from the regulatory pathway to explain the stronger impact of activity-dependent BDNF secretion on synaptic plasticity (Thomas **and Davies 2005**). As long as the value of the second term on the right-hand side of Eq. (35) is positive and negligibly small, omitting the term does not alter the solution of the equation considerably. Consequently, we omit the term and we obtain

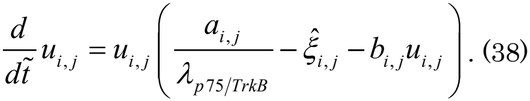

### 2.5 Reduction to single-spine dynamics

Since it is impossible to analytically solve the system of simultaneous equations given by Eq. (38), we can reduce it to the dynamic equation of a single spine by applying the effective medium approximation, in which all spine volumes are assumed to be given by the stationary uniform solution 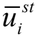 to Eq. (38) as follows:

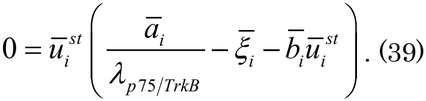

Here, 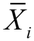 indicates *X*_*i, j*_ averaged over all spine sites. Therefore,

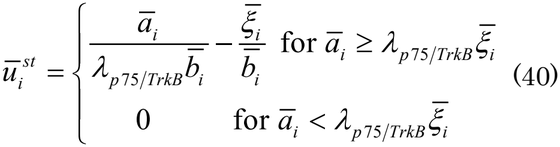

and

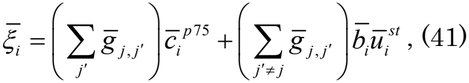

where we replace 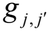 with their average 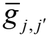 (**Supplementary Information A)** for simplicity. 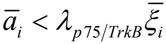 is satisfied only when [*TrkB*] [*p*75 ^*NTR*^], which occurs before the onset of light exposure, that is, eye opening. Thus, within the critical period, we can set 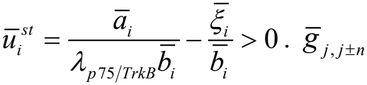 is given by

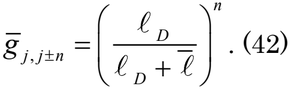

Combining Eqs. (40), (41), and (42), we obtain

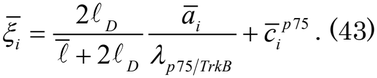

Taken together, by applying the effective medium approximation in which all 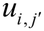 ‘s other than *u*_*i, j*_ are replaced with 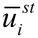, we can reduce Eq. (39) to a set of single variable differential equations:

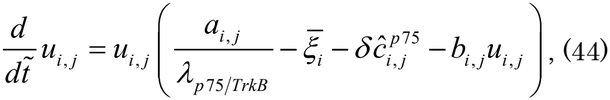

where 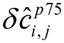 represents the spine-site-dependent fluctuation around 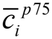 . This equation describes that *u*_*i, j*_ (*t*) approaches the stationary states 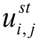, fluctuating within the timescale of changes in *a*_*i, j*_ and *b*_*i, j*_, which is much longer than the timescale of changes in individual spine volumes *u*_*i, j*_ (*t*).

### 2.6 Activity-dependent synaptic rewiring

The generation and elimination of spines are described as the transitions among different stationary state solutions to Eq. (44).

By setting the time derivatives 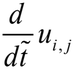 on the left-hand side in Eq. (44) to 0, we can obtain the stationary-state solution 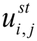 as

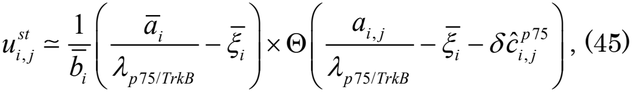

with the Heaviside function Θ (*x*) . This indicates that a spine is generated (eliminated) when 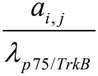 is greater (smaller) than 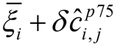 . The generated spine volume is approximately given by 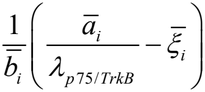 . The transition probability of spine generation/elimination is obtained by averaging the Heaviside function with respect to the stochastic variable 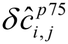 . Here, we assume 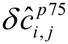 to be the Gaussian random variable with mean 0 and standard deviation 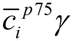 . The transition probability is well approximated by the logistic function (**Supplementary Information B; Tanaka, 1990**).

Generally, the value 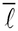 of is unknown. To eliminate it in the model, we simplify the model assuming that the total length of dendrites is preserved while synapses are rewired:

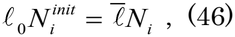

where *l*_0_ and 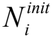 are the initial interspine interval and the initial number of spines on the *i* th cortical neuron, respectively. For further simplicity, a single dendrite is infinitely long and all dendrites belonging to the *i* th neuron are replicas of a single dendrite. Thereby, we can eliminate the average interspine interval 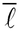 as 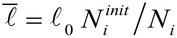.

When we argue the rewiring of LGN afferent synaptic inputs to individual cortical neurons, we assume that all mature spines establish synaptic contacts with one of the boutons on the LGN axons. This means that at every spine site, we can express *a*_*i, j*_ and *b*_*i, j*_ as 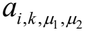 and 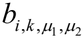, respectively, using the index of the LGN cell (*k, μ*_1_, *μ*_2_), which sends a synaptic input to the spine *j* . A spine connecting with a bouton on the axon of an LGN cell (*k, μ*_1_, *μ*_2_) may change to connect with a bouton on the axon of another LGN cell (*k*′, 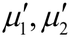) when spine motility is high. Such a spine-bouton contact, namely, a synapse, is updated by the Monte Carlo simulation method (**Metropolis et al., 1953**). For the sake of convenience, one simulation step is defined as 48× 48 (number of model cortical neurons) × 53× 4 (maximum number of model LGN cells that can send afferent inputs to a single cortical neuron) trials of updates of single spines. Hereafter, we denote the simulation step as *t* in the evolution of receptive fields and the related maps.

Let us define 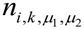 as

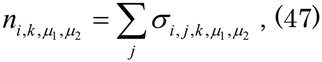

representing the number of afferent synaptic inputs from the LGN cell (*k, μ*_1_, *μ*_2_) to the *i* th cortical neuron. At the transition state between 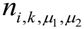 and 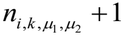, the number of synapses may well be given by 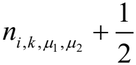, which is the intermediate value between before and after the transition. Using the logistic function form of the transition probability, we respectively write the probabilities of generation and elimination of a synapse as

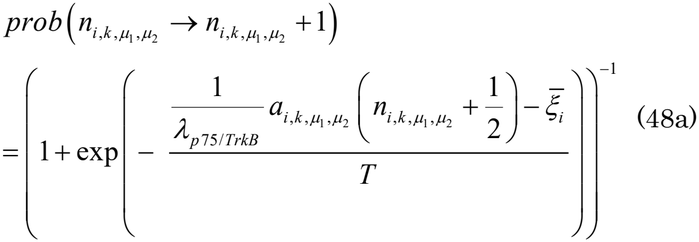

and

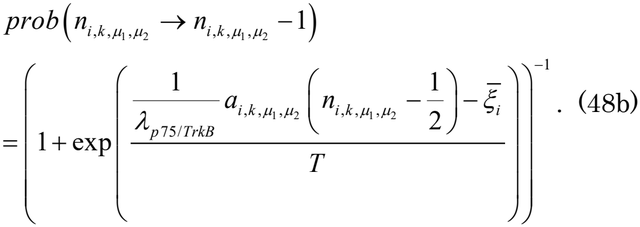

Here, the fictitious temperature *T* is given by

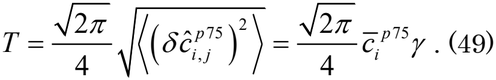

Spontaneous spine elongation and shrinkage even in the absence of external stimuli are considered to be mediated by the polymerization and depolymerization of actin filaments that form the morphology of spines, although the detailed molecular mechanisms of this type of spine motility are not necessarily elucidated in depth. In any case, spine motility may work for the random search of more appropriate axonal boutons to form contacts (**Bonhoeffer and Yuste, 2002**). During the critical period, such spontaneous spine motility is expected to be high. In contrast, after the end of the critical period, spine motility is speculated to be sufficiently low, indicating that the transition probabilities become sufficiently small and that spines are consolidated and the synaptic circuit hardly changes any longer. In computer simulations, the closure of the critical period is represented by the end of the simulation.

The average values of 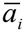 ‘s are calculated every simulation step. The maximum number of afferent synaptic inputs from a single LGN cell to a single cortical neuron is given by *n*_max_.

## 3. Computer simulation

LGN synaptic afferent inputs to individual cortical neurons are updated according to the transition probabilities (48a) and (48b) in the Monte Carlo simulation (**Metropolis et al., 1953**), which describes the generation/elimination of synapses on a timescale from hours to days. The end of the critical period brought about by the reduced spine motility is simply represented by the cessation of the Monte Carlo simulation.

### 3.1 Model LGN and model visual cortex

In the cat LGN, there are lagged and nonlagged cells (**Saul and Humphrey, 1990; 1992**), and spatiotemporal receptive fields of ON-center nonlagged, OFF-center nonlagged, ON-center lagged, and OFF-center lagged cells have been elucidated by the reverse-correlation method of electrophysiology (**DeAngelis et al., 1995**). In this study, we use the mathematical models of the spatiotemporally separable receptive fields of LGN cells that were used in our previous paper (**Tanaka et al., 2020**), and assume that a model LGN is composed of four fictitious layers of the four types of cells for the sake of convenience. Each layer is given as a square lattice of 24×24 cells that are arranged retinotopically. The visual cortex is composed of 48×48 cells which are arranged in a square lattice (**Tanaka et al., 2020**). All parameter values used in the simulations are listed in **Table 1**.

**Table 1.**
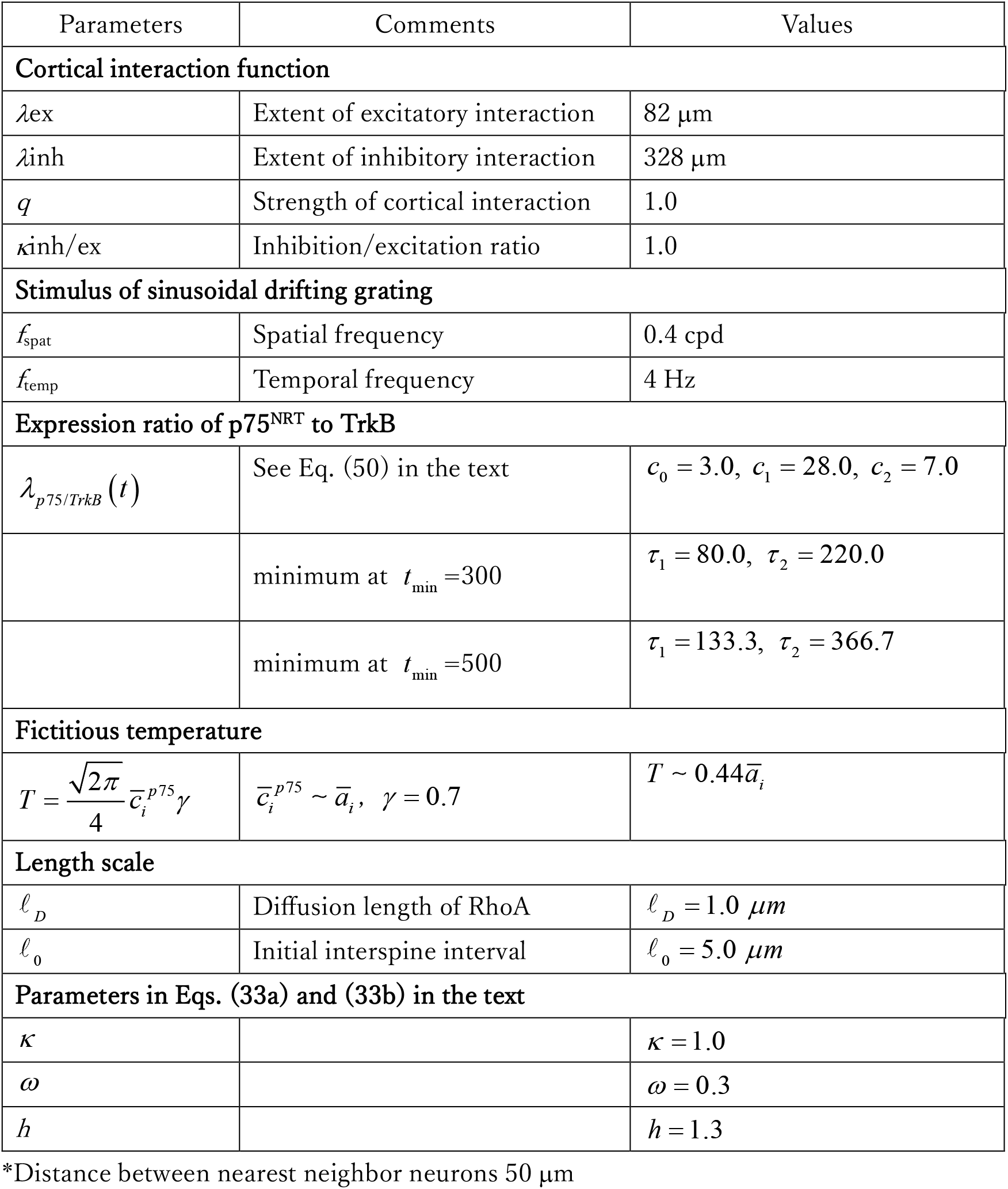
List of parameters and their values used in the simulation.

| Parameters | Comments | Values |
|---|---|---|
| **Cortical interaction function** |  |  |
| *λ*ex | Extent of excitatory interaction | 82 µm |
| *λ*inh | Extent of inhibitory interaction | 328 µm |
| *q* | Strength of cortical interaction | 1.0 |
| *κ*inh/ex | Inhibition/excitation ratio | 1.0 |
| **Stimulus of sinusoidal drifting grating** |  |  |
| *f*<sub>spat</sub> | Spatial frequency | 0.4 cpd |
| *f*<sub>temp</sub> | Temporal frequency | 4 Hz |
| **Expression ratio of p75<sup>NRT</sup> to TrkB** |  |  |
| $\lambda_{p75/TrkB}\left(t\right)$ | See Eq. (50) in the text | $c_0 = 3.0,\ c_1 = 28.0,\ c_2 = 7.0$ |
|  | minimum at $t_{\min}$ =300 | $\tau_1 = 80.0,\ \tau_2 = 220.0$ |
|  | minimum at $t_{\min}$ =500 | $\tau_1 = 133.3,\ \tau_2 = 366.7$ |
| **Fictitious temperature** |  |  |
| $T = \frac{\sqrt{2\pi}}{4}\bar{c}_i^{p75}\gamma$ | $\bar{c}_i^{p75} \sim \bar{a}_i,\ \ \gamma = 0.7$ | $T \sim 0.44\bar{a}_i$ |
| **Length scale** |  |  |
| $\ell_D$ | Diffusion length of RhoA | $\ell_D = 1.0\ \mu m$ |
| $\ell_0$ | Initial interspine interval | $\ell_0 = 5.0\ \mu m$ |
| **Parameters in Eqs. (33a) and (33b) in the text** |  |  |
| *κ* |  | *κ* = 1.0 |
| *ω* |  | *ω* = 0.3 |
| *h* |  | *h* = 1.3 |

\*Distance between nearest neighbor neurons 50 µm

### 3.2 Visual experience

The membrane potential of model LGN cells in response to visual stimulation was calculated by the convolution of visual stimuli with LGN receptive fields. The visual stimuli were given by sinusoidal drifting gratings (**Table 1**). The firing rate of LGN cells, 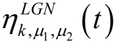, was calculated by the half-rectification of the membrane potential of the cells. It is considered that LGN cells respond most strongly to visual stimuli with spatiotemporal frequencies characteristic of their receptive fields among various stimuli. Hence, to save time for simulations, we presented randomly grating stimuli drifting in 12 directions with spatiotemporal frequencies that were almost the same as the LGN characteristic frequencies (**Movshon et al., 1978; Tanaka et al., 2020**).

### 3.3 Ratio of expression level of p75^NTR^ to that of TrkB

To discuss the developmental changes in the number of synaptic connections and map formation under the changes in the amounts of TrkB and p75^NTR^, we use the time-dependent ratio *λ*_*p* 75/*TrkB*_ . Upon eye opening, the mRNA expression level of p75^NTR^, and hence, the p75^NTR^ protein level begin to decrease (**Bracken and Turrigiano, 2009**), and then TrkB is gradually dephosphorylated by PTPσ to lose its function partially (**Kurihara and Yamashita, 2012**). In contrast, the amount of TrkB does not change considerably throughout postnatal development (**Bracken and Turrigiano, 2009**). Thus, the ratio *λ*_*p* 75/*TrkB*_ decreases rapidly immediately after eye opening and then increases to some extent. This leads to the time dependence of *λ*_*p* 75/*TrkB*_, which can be modeled as the following expression:

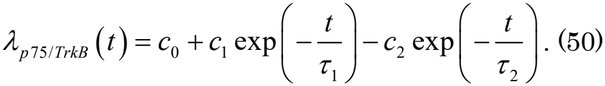

### 3.4 Definition of orientation and direction selectivity

The response of a single neuron is defined as follows: When grating patterns drifting in the direction *θ* orthogonal to the grating orientation *φ* are presented to the model visual field, the response is represented as 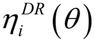 . The response to the grating patterns at the orientation *φ* is represented as 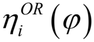 . The relationship between the orientational response and the directional one is given by

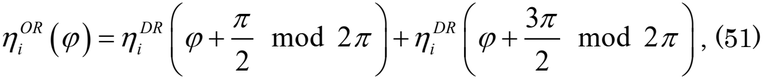

where *x* mod 2*π* indicates a remainder of *x*, when *x* is divided by 2*π*.

The orientation selectivity index (OSI) and direction selectivity index (DSI) of the *i* th cortical neuron are calculated as

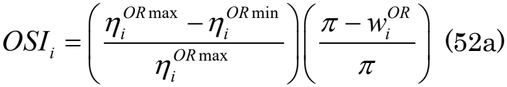

and

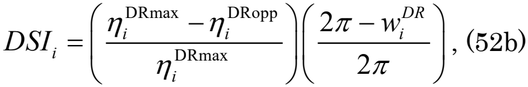

where 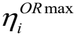and 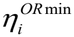 represent the maximum response at the preferred orientation and the minimum response of the *i* th neuron in the range of [0,*π*), and 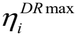 and 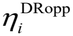 represent, respectively, the maximum response at the preferred direction and the response opposite to the preferred direction of the *i* th neuron in the range of [0, 2*π*), respectively. 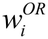 and 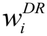 indicate the full widths at half maximum of orientation and direction tuning curves, respectively. The selectivity indices averaged over all cortical neurons are given by

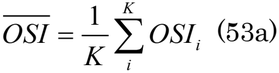

and

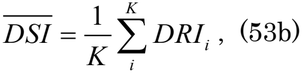

where *K* indicates the total number of cortical neurons *K* (= 48× 48).

## 4. Results

### 4.1 Orientation and direction map development

Figure 1 shows two cases of *λ*_*p 75/TrkB*_, which give a minimum at the simulation step *t* = 300, or 500. Using *λ*_*p 75/TrkB*_, which gives a minimum at *t* = 300, we carried out the simulation until *t* = 2000 when the maps were almost unchanged. Figure 2 shows the orientation maps (Fig. 2a) and direction maps (Fig. 2b) at *t* = 100, 300, 500, 1000, and 2000. The color in the maps indicates preferred orientations (Fig. 2a) and preferred directions (Fig. 2b). The brightness represents orientation and direction selectivity in the respective maps. At *t* = 100, preferred orientations and directions were randomly arranged without organized map structures. At *t* = 300, iso-orientation domains were clearly visible, although they were not necessarily sharply delineated. In contrast, iso-direction domains were scarcely visible, and a large portion of neurons exhibited weak direction selectivity because most pixels were drawn in the dark. From *t* = 300 to 2000 steps, the basic structure of the orientation map was well preserved, whereas the borders of iso-orientation domains became gradually sharper. On the other hand, iso-direction domains were expanding until *t* = 2000, but dark pixels indicating neurons with low direction selectivity formed fractures. In this simulation, the Monte Carlo update is stopped at *t* = 2000, which indicates that the critical period closes around *t* = 2000.

**Figure 1.**
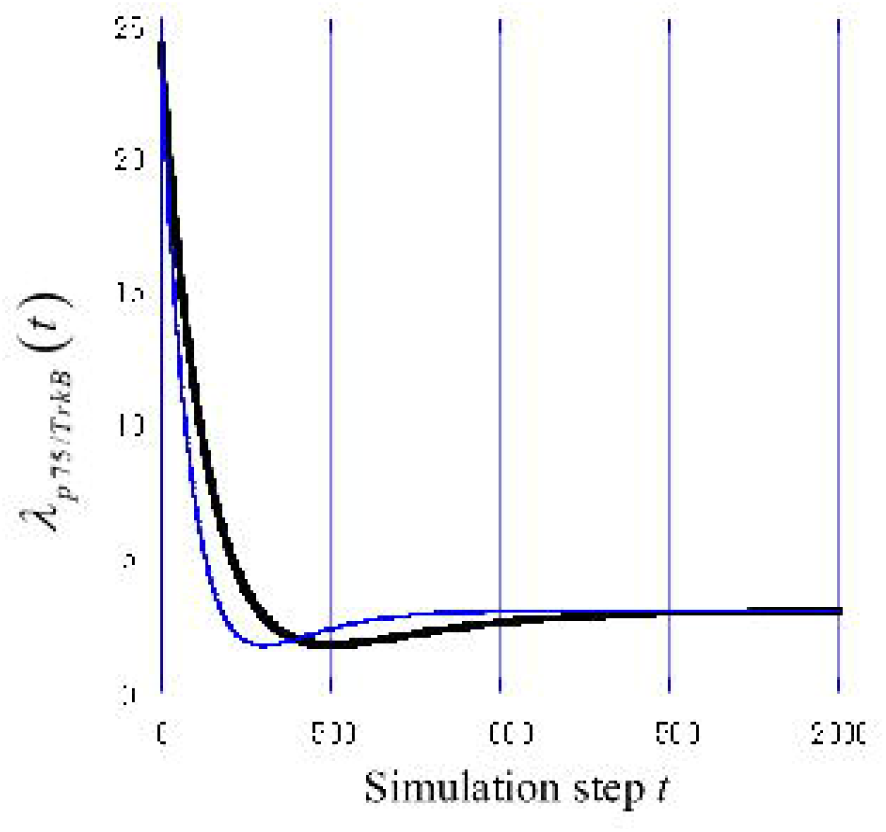
Simulation-step dependence of *λ*_*p* 75/*TrkB*_ (*t*) The blue and black curves indicate *λ*_*p* 75/*TrkB*_ (*t*) that give a minimum at *t*_min_ = 300 and 500, respectively.

**Figure 2.**
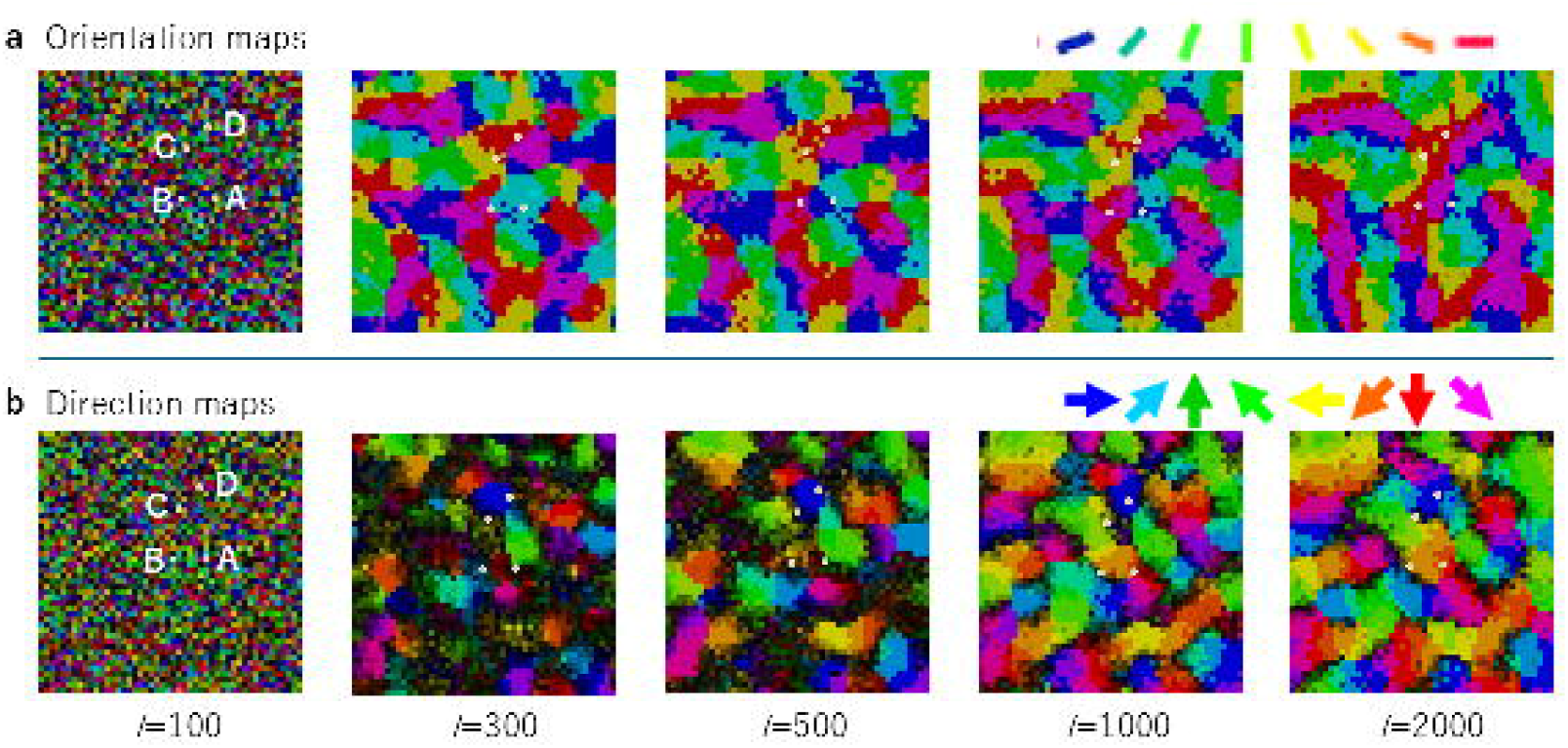
Orientation and direction map development. Orientation maps (Fig. 2a) and direction maps (Fig. 2b) at *t* = 100, 300, 500, 1000, and 2000. The color in the maps indicates preferred orientations and preferred directions. The brightness represents orientation and direction selectivity in the respective maps. At *t* = 100, preferred orientations and directions were randomly arranged without organized map structures. At *t* = 300, iso-orientation domains were clearly visible, although they were not necessarily sharply delineated. In contrast, iso-direction domains were scarcely visible, and a large portion of neurons exhibited very weak direction selectivity. From *t* = 300 to 2000 steps, the borders of iso-orientation domains were gradually sharper, but the basic structure was well preserved. On the other hand, iso-direction domains were expanding until *t* = 2000, but dark pixels indicating neurons with low direction selectivity formed fractures.

It is important that this model well reproduced pinwheel-like arrangement of preferred orientation around orientation point singularities as observed in the optical imaging experiment (**Bonhoeffer and Grinvald, 1991**) and direction reversal at low direction-selective regions as reported in the experiment (**Shmuel and Grinvald, 1996**).

### 4.2 Direction tuning curves

Figure 3 shows the direction tuning curves of four typical cortical neurons at *t* = 100, 300, 500, 1000, 1500, and 2000. They were taken from the neurons located at A (30, 24), B (24, 24), C (25, 33), and D(29,37), as indicated by small white dots in orientation and direction maps (Fig. 2). Neurons A and B in Fig. 3 show changes from bimodal curves to unimodal curves, indicating that response selectivity changes from bi-direction selectivity (low direction selectivity) to uni-direction selectivity (high direction selectivity). This behavior indicates that orientation selectivity is formed first and direction selectivity is formed later in these neurons. Neuron C depicted in Fig. 3 maintains response selectivity to be bi-directional from *t* = 300 to 2000, because it is located on the direction fracture. Neuron D in Fig. 3 shows high direction selectivity throughout from *t* = 300 to 2000.

**Figure 3.**
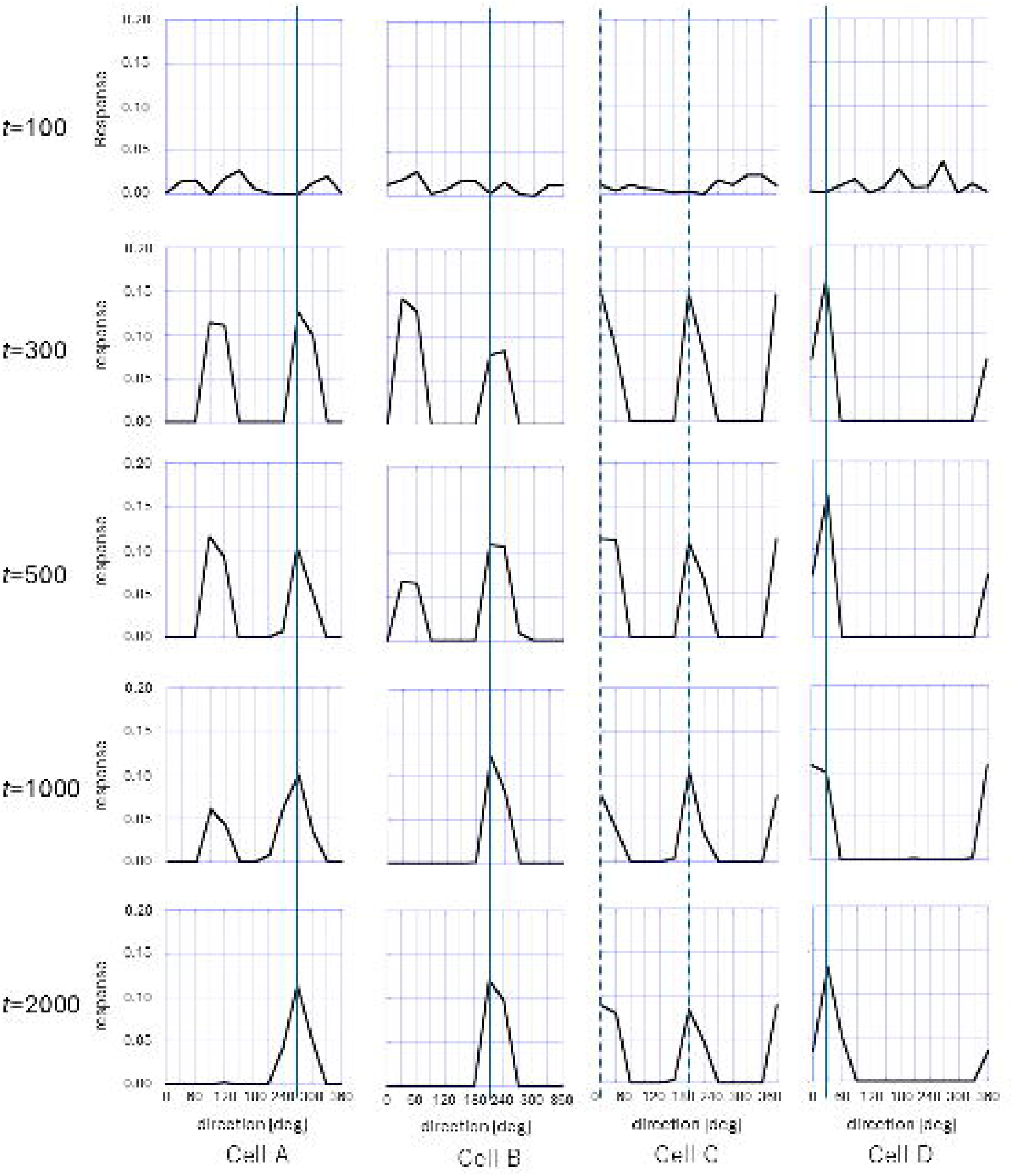
Orientation and direction tuning curves. Direction tuning curves of four typical cortical neurons at *t* = 100, 300, 500, 1000, and 2000 are shown. They were taken from the neurons located at A (30, 24), B (24, 24), C (25, 33), and D(29, 37), as indicated by small white dots in orientation and direction maps (Fig. 2).

OSI and DSI of neuron A – D are shown in Fig. 4. In all four figures, OSI (blue curves) reached maximum values around *t* = 300 and remained high up to *t* = 2000, although DSI increases (green curves) were different in the four figures. Neuron A showed a gradual increase in direction selectivity. Neuron B showed a rapid increase in DSI, but it began to increase later than the increase in orientation selectivity. The DSI of neuron C was low throughout all the simulation steps, which is consistent with the bi-directional behavior of the tuning curves of neuron C shown in Fig. 3. Neuron D showed high DSI from the early steps, because it was located in the middle of the iso-direction domain already formed at *t* = 300.

**Figure 4.**
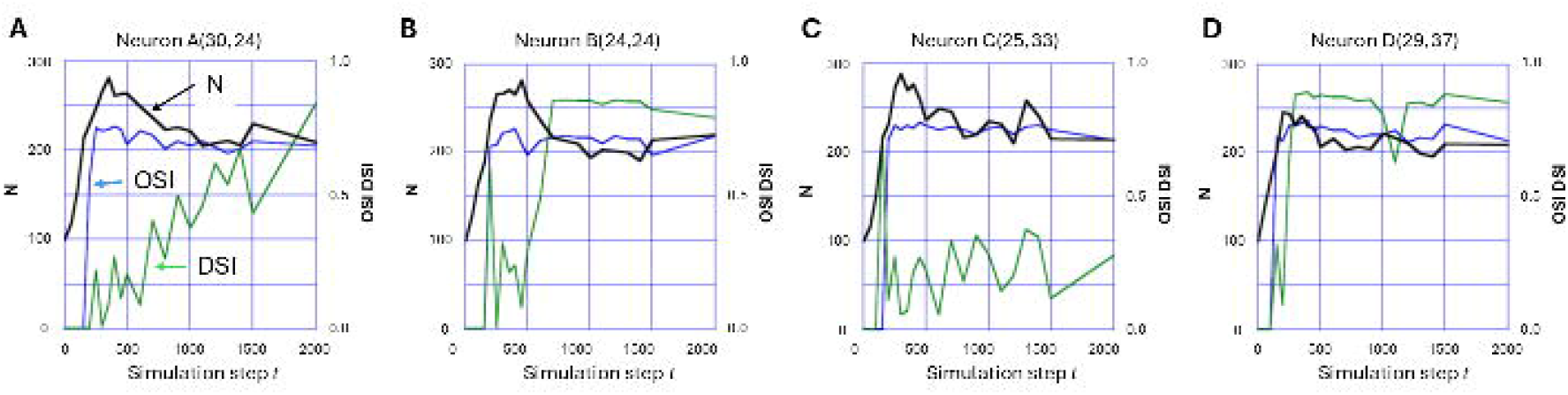
Changes in OSI, DSI and *N* of four typical neurons. OSI (blue curves) reached maximum values around *t* = 300 and remained high up to *t* = 2000, although DSI increases (green curves) were different in the four neurons. Neuron A showed a gradual increase in direction selectivity. Neuron B showed a rapid increase in DSI, but it began to increase later than the maturation of orientation selectivity. The DSI of neuron C was low throughout all the simulation steps, which is consistent with Fig. 3C. Neuron D showed high DSI from the early steps, because it was located in the middle of the iso-direction domain already formed at *t* = 300. The number of LGN afferent synaptic inputs to cortical neurons (black curves) reached the maximum around the simulation step *t*_min_ = 300 where *λ*_*p 75/TrkB*_ gives a minimum.

### 4.3 Number of LGN afferent inputs

Figure 4 also shows changes in the number of LGN afferent synaptic inputs *N*_*i*_ to the *I* th cortical neuron (black curves) with the simulation step, because we assume that each mature spine has a single synaptic input and that the number of spines is the same as the number of geniculo-cortical synaptic inputs. Compared with Fig. 1, *N*_*i*_ reached the maximum around the simulation step *t*_min_ = 300 where *λ*_*p* 75 /*TrkB*_ gave a minimum.

In all four figures, OSI rapidly increased from the onset of visual experience and reached a maximum value around *t* = 300, where the number of LGN inputs became maximum. On the other hand, it is likely that DSI developed with a delay after the formation of orientation selectivity. Neurons whose direction selectivity was low were clustered to form direction fractures where neurons were bi-directional.

### 4.4 Spatiotemporal receptive fields

Figure 5 shows the spatiotemporal receptive fields of neuron A indicated in Fig. 2, changing with the simulation step *t* . Figure 5a depicts the development of spatial receptive field snapshots at 50 ms after the onset of visual stimulation. ON and OFF subfields were randomly arranged at *t* = 100. At *t* = 300 and later, ON and OFF subfields were segregated vertically and elongated horizontally. This indicates that oriented simple-cell-like receptive fields were formed as early as *t* = 300, and that the neuron was selective to horizontal orientation. Figure 5b shows the development of the x-t plot of receptive fields of the same neuron A. Subfields were elongated vertically but the positions of ON and OFF subfields were exchanged at a certain time until *t* = 1000, which indicates that the receptive field was spatiotemporally separable and non-direction-selective, that is, bi-directional. At *t* = 2000, ON and OFF subfields were elongated obliquely, which indicates that the receptive field was spatiotemporally inseparable. That is, this neuron became direction-selective because the temporal sum occurred when a grating stimulus moved with the velocity given by the inverse of inclination of elongated ON and OFF subfields. Therefore, the neuron became orientation-selective at an early stage of development and direction-selective at a late stage.

**Figure 5.**
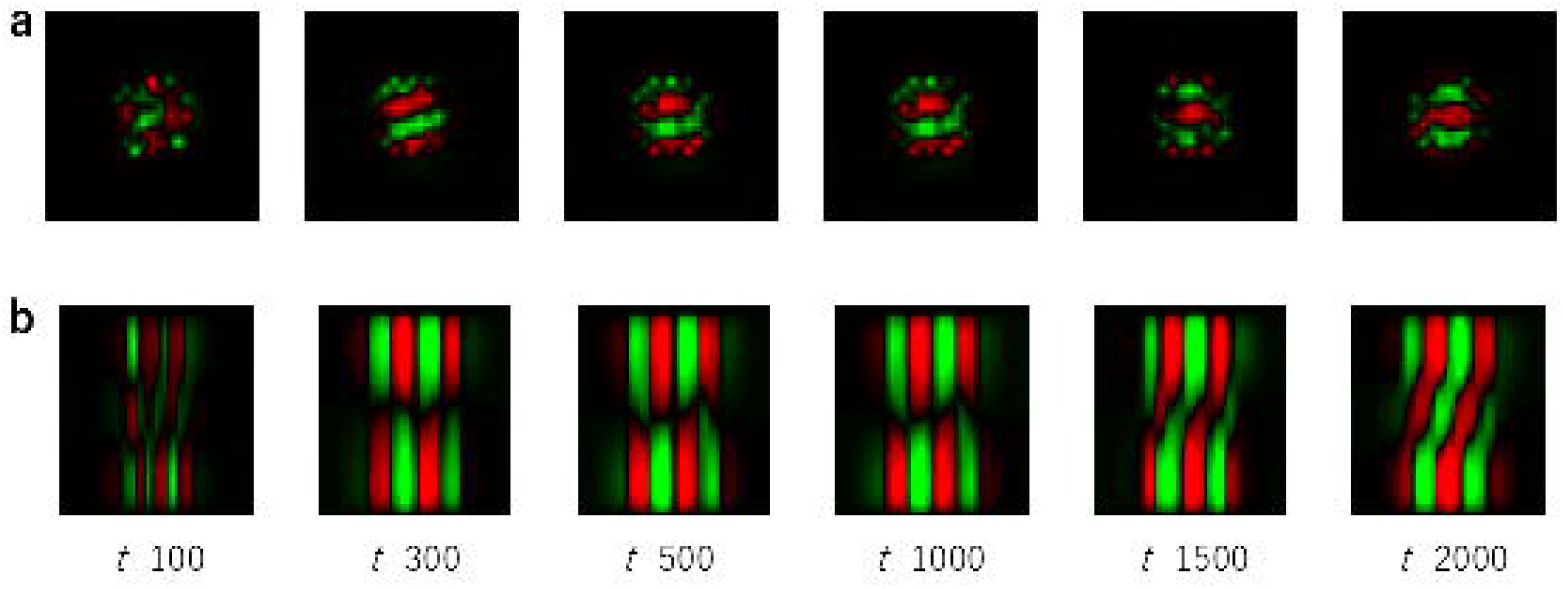
Spatiotemporal receptive fields of neuron A. **a**. Changes of spatial receptive field snapshots at 50 ms after the onset of visual stimulation. At *t* = 300 and later, ON and OFF subfields were elongated horizontally. **b**. ON and OFF subfields were randomly arranged at *t* = 100. Around *t* = 1000, the receptive field changed from spatiotemporally separable to inseparable receptive fields, which is indicative of the emergence of direction selectivity around *t* = 1000.

### 4.5 Distributions of OSI and DSI

Figure 6 shows the distributions of OSI and DSI at *t* = 100, 300, 1000, and 2000. At *t* = 100, most neurons exhibited low OSI and DSI. At *t* = 300, OSI was distributed sharply around 0.7, whereas DSI was widely distributed and biased to lower values. From *t* = 1000 to 2000, most neurons were highly orientation-selective and a large portion of the neurons exhibited high direction selectivity, but low-direction-selective neurons remained, which were located in direction fractures. Direction selectivity increased with the decrease in the LGN synaptic input number, which suggests that synaptic inputs inappropriate for direction selectivity were pruned, whereas orientation selectivity was almost preserved. This situation is consistent with the observations on orientation and direction selectivity formation in our former self-organization model, which constrains the afferent synaptic number in a single cortical neuron (**Tanaka and Miyashita, 2009**).

**Figure 6.**
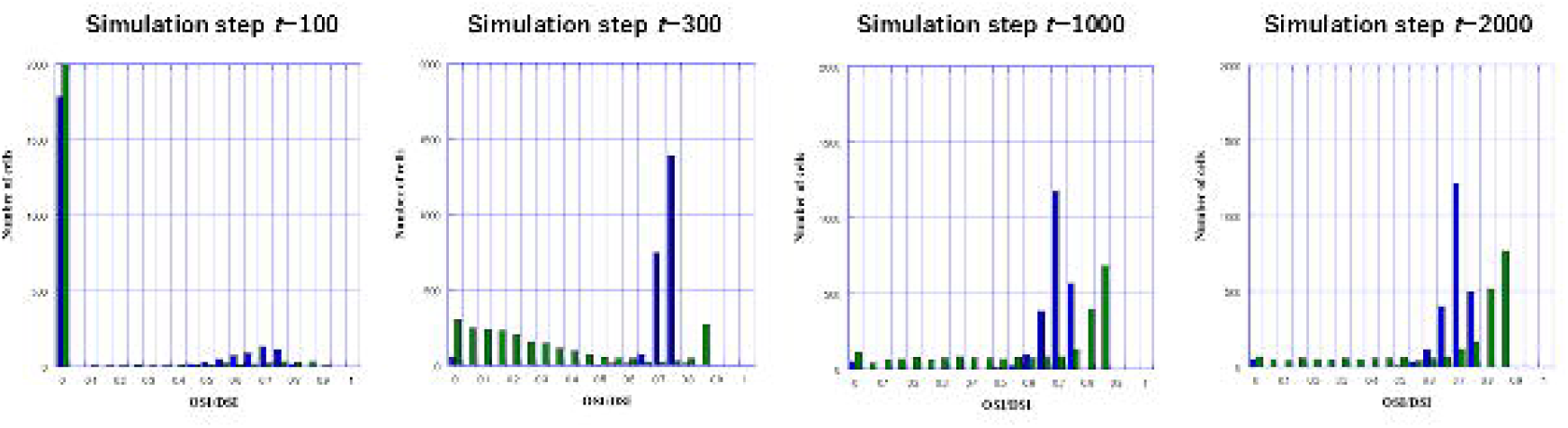
Distributions of OSI and DSI at *t* = 100, 300, 1000, and 2000. At *t* = 100, most neurons exhibited low OSI and DSI. At *t* = 300, OSI was distributed sharply around 0.7, and direction selectivity was widely distributed and biased to lower selectivity. From *t* = 1000 to 2000, most neurons were highly orientation-selective and a large portion of the neurons exhibited high direction selectivity.

### 4.6 Patterns of LGN afferent synaptic inputs

For LGN synaptic inputs to individual cortical neurons, receptive field centers of LGN cells of each type were clustered to form subfields of visual cortical neurons at *t* = 300, as shown in Figs. 7A1 and 7B1. The receptive field arrangements of ON- (OFF-) center nonlagged cells are somehow similar to those of OFF- (ON-) center lagged cells. This is because both ON- (OFF-) center nonlagged cells and OFF- (ON-) center lagged cells have early ON (OFF) and late OFF (ON) responses on the time axis. Such similarity in the temporal response pattern suppresses the separation of subfields. Therefore, ON-center nonlagged (lagged) cells and OFF-center lagged (nonlagged) cells were largely overlapped in the visual field (rightmost panels in Figs. 7A2 and 7B2). These situations are more precisely described by estimating the Hebbian coincidence 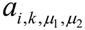 . However, the Hebbian coincidence includes nonlinearity regarding the LGN cell activities, which makes it difficult to estimate analytically. Thus, it is convenient to linearize the half-rectification of the LGN cell activities, as shown in **Supplementary Information C**. The Hebbian coincidence is approximately given by the overlap integral between the two LGN cell activities. As a further simplification, when we assume that visual stimuli are Gaussian white noise, the overlap integral is given by the covariance of two LGN cells’ receptive fields. This indicates that as two LGN receptive fields become more similar, the Hebbian coincidence becomes larger (**Supplementary Information C**).

**Figure 7.**
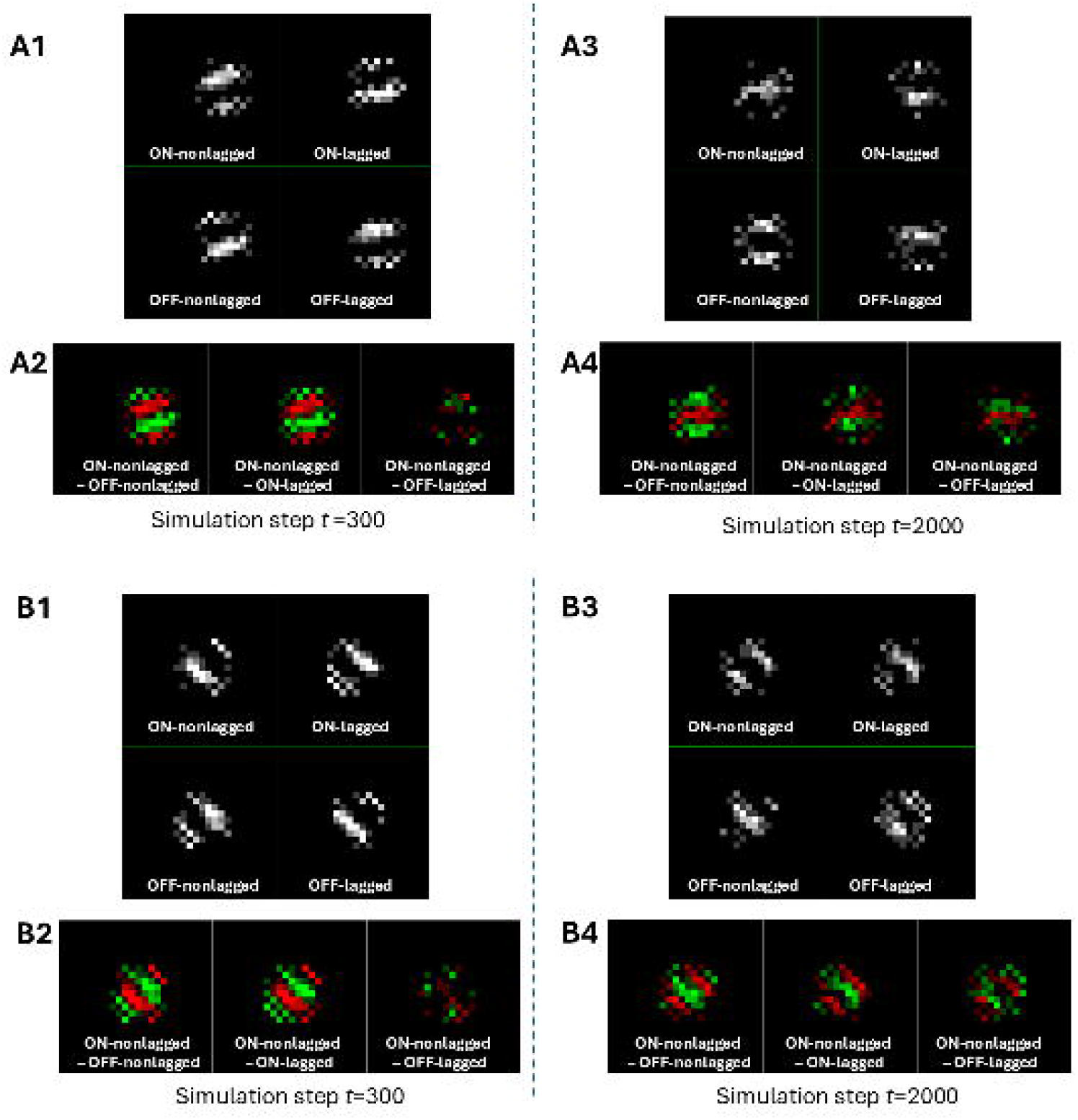
Patterns of LGN afferent synaptic inputs. For LGN synaptic inputs to individual cortical neurons, receptive field centers of LGN cells of each type were clustered to form subfields of visual cortical neurons at *t* = 300, as shown in Figs. 7A1 and 7B1. The receptive field arrangements of ON-nonlagged (OFF-nonlagged) cells are somehow similar to those of OFF-lagged (ON-lagged) cells. This is because both ON-nonlagged (OFF-nonlagged) cells and OFF-lagged (ON-lagged) cells have early ON (OFF) response and late OFF (ON) response in the time axis. Such similarity in the temporal response pattern suppresses the separation of subfields. In contrast, ON-nonlagged (ON-lagged) cells and OFF-nonlagged (OFF-lagged) cells have ON/OFF opposite subfields at the same retinotopic positions. Because of this dissimilarity in spatiotemporal receptive fields, ON-nonlagged (ON-lagged) cells and OFF-lagged (OFF-lagged) cells almost overlapped. However, long-run simulation gradually separates subfields of ON-nonlagged (OFF-nonlagged) cells from those of OFF lagged (ON-lagged) cells. As a result, lagged and nonlagged cells that send inputs to individual cortical neurons were segregated in the visual field to form spatiotemporally inseparable receptive fields, which produced direction-selective neurons in the visual cortex.

Long-run simulation gradually separates subfields of ON- (OFF-) center nonlagged cells from those of OFF- (ON-) center lagged cells (Figs. 7A3 and 7B3). As a result, lagged and nonlagged cells that send inputs to individual cortical neurons were segregated in the visual field (rightmost panels in Figs. 7A4 and 7B4) to form spatiotemporally inseparable receptive fields, which produced direction-selective neurons in the visual cortex. In contrast, ON-center nonlagged (lagged) cells and OFF-center nonlagged (lagged) cells have ON/OFF opposite polarities, which promotes the segregation of ON- and OFF-center nonlagged cell inputs throughout all the simulation steps, forming orientation-selective receptive fields (leftmost panels in Figs. 7A2, 7A4, 7B2 and 7B4). As for lagged cells, the situation is the same as that for nonlagged cells (data not shown). Thus, the formation of orientation-selective and direction-selective receptive fields takes place as the rearrangement of LGN receptive fields in a spontaneous symmetry breaking manner.

### 4.7 Developmental profiles of average selectivity

Figure 8 shows the developmental profiles of OSI, DSI and the number of geniculo-cortical connections averaged over all model cortical neurons for the two cases where *λ*_*p 75/TrkB*_ gives a minimum at *t* = 300 (a1) or 500 (b1). The number of geniculo-cortical connections (black curve) monotonically increased and took a maximum around *t* = 300 (a1) or 500 (b1), and then decreased. After *t* = 1500, it became a constant at about 220. OSI (blue curve) rapidly increased immediately after the onset of visual stimulation at *t* = 0 and reached a maximum at *t* = 300 (a1) or 500 (b1). Then it decreased slightly and asymptotically took a constant value around OSI = 0.70. DSI (green curve) also increased after the onset of stimulation up to *t* = 300 (a1) or 500 (b1) and then restarted to increase until *t* = 1500 (a1 and b1), reaching a plateau around DSI = 0.67. The maturation of direction selectivity was clearly delayed later than that of orientation selectivity, which agrees well with experimental observations (Li **et al., 2006; White et al., 2007**). Figures 8a2 (b2) and 8a3 (b3) show orientation maps and direction maps at *t* = 2000, respectively. Comparison of Figs. 8a2 (a3) and Fig. 8b2 (b3) shows that the basic structures of those maps were unaffected by whether *λ*_*p 75/TrkB*_ took a minimum at *t* = 300 or 500.

**Figure 8.**
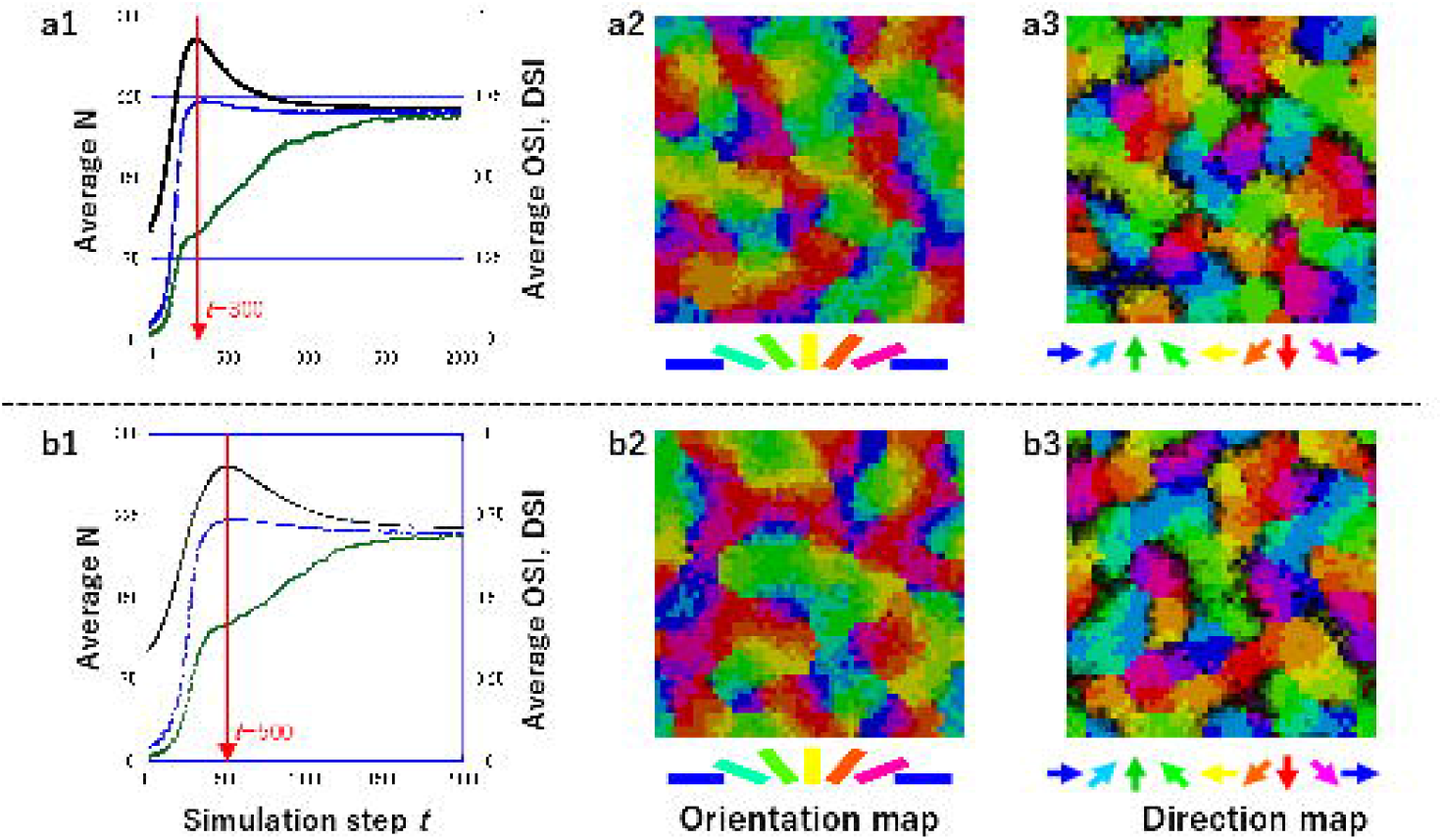
Developmental profiles of OSI, DSI, and *N* averaged over all model cortical neurons. The two cases where *λ*_*p 75/TrkB*_ gives a minimum at *t* = 300 (a1) and 500 (b1) are shown. *N* monotonically increased (black curve) and took a maximum around *t* = 300 (a1) or 500 (b1) and then decreased. After *t* = 1500, it became a constant at about 220. OSI (blue curve) rapidly increased immediately after the onset of visual stimulation at *t* = 0 and reached a maximum at *t* = 300 (a1) or 500 (b1). Then it decreased slightly and asymptotically reached a constant value around OSI = 0.70. DSI (green curve) also increased after the onset of stimulation up to *t* = 300 (a1) or 500 (b1) and then restarted to increase until *t* = 1500 (a1 and b1), reaching a plateau around DSI = 0.67. The maturation of direction selectivity was delayed behind that of orientation selectivity, which agrees well with experimental observations. Figures 8a2 (b2) and 8a3 (b3) show orientation maps and direction maps at *t* = 2000, respectively. Comparison of Figs. 8a2 (a3) and 8b2 (b3) shows that the basic structures of those maps were unaffected by whether *λ*_*p 75/TrkB*_ took a minimum at *t* = 300 or 500.

Furthermore, we performed simulations for different initial patterns of geniculo-cortical inputs and different random sequences for Monte Carlo simulation with the same number of initial connections *N*_*init*_ = 100. Despite such randomness, curves for OSI, DSI, and the number of synapses were almost the same (not shown).

## 5. Discussion

### 5.1 Development of orientation and direction maps

Iso-orientation domains are clearly delineated at *t* = 300 when OSI reaches its peak (Figs. 8a1 and 8b1), and the orientation map does not change considerably afterward (Fig. 2a). In contrast, some regions exhibiting low DSI (dark domains) remain until *t* = 500 (Fig. 2b). Neurons with low DSI exhibit bi-directional responses to drifting grating stimuli, as shown in tuning curves of neurons A, B, and C in Fig. 3. The bimodal tuning curves of neurons A and B before *t* = 1000 change to unimodal tuning curves by *t* = 2000. At *t* = 1000 or later, iso-direction domains spread almost over the model visual cortex.

However, narrow regions of low DSI, which are often called direction fractures, are exceptionally preserved even at *t* = 2000 (Fig. 2b). Along the direction fractures, neurons are bi-directional, as seen in neuron C (Fig. 3). The observation that direction selectivity is formed sufficiently later than the emergence of orientation selectivity agrees with experimental findings (**Li et al., 2006; White and Fitzpatrick, 2007**).

### 5.2 Development of spatiotemporal receptive fields

Figure 5 shows developmental changes in x-t plots of spatiotemporal receptive fields at a cortical neuron located at A(30, 24) in the simulation performed using *λ*_*p 75/TrkB*_ giving a minimum at *t* =300. We can see the transition from spatiotemporally separable receptive fields to inseparable receptive fields around *t* =1500. Direction maps are likely to mature with the reduction in the total number of synaptic inputs from the LGN in the late phase.

In simulations performed with strong intracortical interactions, iso-direction domains tend to be delineated because neurons of opposite preferred directions are likely to be segregated in the cortex, whereas they are allowed to be closely located, and thus direction maps tend to be randomly organized in simulations with weak intracortical interactions (**Supplementary Information D**). However, when afferent synaptic inputs are abundant, strong dissimilarity between LGN receptive field types facilitates the formation of orientation selectivity. In contrast, as the number of afferent inputs decreases, weak dissimilarity between ON (OFF)-center nonlagged and OFF (ON)-center lagged receptive fields facilitates the segregation of afferent inputs to form spatiotemporal receptive fields, resulting in the formation of direction selectivity, as shown in Fig. 7.

### 5.3 Developmental changes in N, OSI, and DSI

Figure 8 indicates that (1) *N* becomes maximum at *t*_min_ = 300 or 500 at which *λ*_*p 75/TrkB*_ takes a minimum value; (2) orientation selectivity is formed in a relatively early stage of development, takes a maximum value at *t*_min_ = 300 or 500, and then slightly decreases; and (3) direction selectivity also increases from *t* = 0, reaches a constant value at *t*_min_ = 300 or 500, and then increases again up to a plateau. These observations again reflect that direction selectivity develops after orientation selectivity formation matures (**Li et al., 2006; White and Fitzpatrick, 2007**). While synaptic inputs from the LGN are eliminated after *t*_min_ = 300 or 500, direction selectivity is enhanced. This suggests that the pruning of inappropriate synaptic connections shapes direction selectivity more effectively than orientation selectivity.

### 5.4 About the critical period from viewpoints of this model

This model of visual cortical development was built mainly on the basis of the BDNF effects and developmental changes in the TrkB and p75^NTR^ expression levels. BDNF released both in activity-dependent and activity-independent manners. The former pathway contributes to the generation of appropriate dendritic spines and the elimination of inappropriate spines, whereas the latter is assumed to bring about fluctuations in spine motility. Since the binding of mature BDNF with TrkB receptors generates dendritic spines, the decrease in the p75^NTR^ expression level after the onset of light experience may start the critical period of the visual cortex. On the other hand, CSPGs bind with PTPσ, which dephospholyrates active TrkB. The increased expression level of inactive TrkB in turn decreases the amount of Cdc42 and Rac1 and suppresses spine elongation and maturation. Furthermore, CSPGs may directly block spine motility (**de Vivo et al, 2013**). As a result, the increase in the amount of CSPGs weakens synaptic plasticity and finally leads to the closure of the critical period. Namely, in this model, a major mechanism of synaptic rewiring is based on the activation of Rho-GTPases caused by the binding between BDNF and TrkB, although there would be other molecular mechanisms contributing to learning via long-term potentiation and depression after the closure of the critical period. In this study, we omitted the phenomenon of the fading out of the critical period under the effects of CSPGs in order to focus on synaptic rewiring during the critical period. In the future study, we will take into account changes in spine motility in the extracellular matrix containing CSPGs to describe how the critical period closes.

This model is advantageous over our previous model in which the number of synaptic inputs from the LGN is constant (**Tanaka et al., 2020**) because we can account for the developmental changes in the number of synaptic inputs. Moreover, this model is unique because it links molecular mechanisms to orientation/direction selectivity and map formation, which cannot be explained by computational theories for map self-organization (**Durbin and Mitchison, 1990; Obermayer et al., 1990; Miller, 1994; Carreira-Perpinã′n, 2005**).

## Acknowledgement

This work was supported by JSPS KAKENHI Grant Number 19K12743 and Takahashi Industrial and Economic Research Foundation.

